# Decoding DNA sequence-driven evolution of the human brain epigenome at cellular resolution

**DOI:** 10.1101/2023.09.14.557820

**Authors:** Emre Caglayan, Genevieve Konopka

**Affiliations:** Department of Neuroscience, UT Southwestern Medical Center, Dallas, TX 75390, USA; Peter O’Donnell Jr. Brain Institute, UT Southwestern Medical Center, Dallas, TX 75390, USA

## Abstract

DNA-based evolutionary comparisons of regulatory genomic elements enable insight into functional changes, overcoming tissue inaccessibility. Here, we harnessed adult and fetal cortex single-cell ATAC-seq datasets to uncover DNA substitutions specific to the human and human-ancestral lineages within apes. We found that fetal microglia identity is evolutionarily divergent in all lineages, whereas other cell types are conserved. Using multiomic datasets, we further identified genes linked to multiple lineage-divergent gene regulatory elements and implicated biological pathways associated with these divergent features. We also uncovered patterns of transcription factor binding site evolution across lineages and identified expansion of bHLH-PAS factor targets in human-hominin lineages, and MEF2 factor targets in the ape lineage. Finally, conserved features were more enriched in brain disease variants, whereas there was no distinct enrichment on the human lineage compared to its ancestral lineages. Our study identifies major evolutionary patterns in the human brain epigenome at cellular resolution.

## Introduction

Comparative genomics is widely used to investigate genomic evolutionary patterns^1,2^. Genomic novelties in the human lineage are exceptionally well studied due to their importance in understanding human evolution. Notable examples include human accelerated regions (HARs) that possess a significantly greater number of DNA substitutions in the human lineage compared to chimpanzees and other non-human species^3^, human-specific deletions in otherwise conserved regions (hCONDELs)^4,5^, and newly emerged human-specific regulatory elements (HAQERs)^6^. Functional mechanisms of these elements are investigated both per candidate region^7,8^ and across all candidate regions using functional omics strategies^9–11^. More recent studies have also focused on evolutionary changes in the ancestral lineages of humans that offer additional insight into the composition of the human genome^12,13^.

In addition to exploring the newly evolved features of the human genome, functional aspects of genomic features (e.g., genes and gene regulatory elements) can also be comparatively analyzed in a high-throughput manner. Single-cell transcriptomic and epigenomic strategies are particularly powerful for understanding gene and gene regulatory element (GRE) activity since the molecular landscape of the human brain contains various specialized cell types (e.g., neurons, oligodendrocytes) and numerous cellular subtypes (e.g., upper layer excitatory neurons)^14^. Indeed, single-cell technologies have been applied to human and non-human primate brains to uncover human-specific molecular changes at cell type resolution^15–18^. However, these studies are constrained by the availability of high-quality brain tissue from non-human primates. Such tissues can be scarce from endangered species, especially from developmental periods. Most studies to date have only compared human brain to chimpanzee brain among the ape species and have focused mainly on the adult brain tissues^15–18^. One potential solution for gaining evolutionary insights is to utilize cellularly resolved epigenomes of the human brain to identify putative GREs and explore their sequence divergence among ape species.

Here, we implemented this solution through an integrative analysis of single-cell ATAC-seq and multi-omics datasets from adult and fetal human cortex. Focusing on the evolution of the human brain epigenome within the ape lineage, we identified unique substitutions per GRE in humans and their ancestral lineages by comparing DNA sequences across apes, old world monkeys and new world monkeys. We found unexpected cell type evolution patterns in fetal and adult brains, utilized multiomic datasets to identify genes with a significant excess of divergent GREs per lineage, uncovered transcription factor binding site (TFBS) expansion patterns, and investigated disease susceptibility of divergent and conserved GREs. Our results uncover major patterns and provide a comprehensive overview of the human brain epigenome evolution.

## Results

### Identification of lineage specific substitutions

We utilized single-nuclei ATAC-seq datasets from fetal (post-conceptional weeks 16-24) and adult human brain cortex^16,19^ for a comprehensive list of putative GREs **(Supplemental Figure 1a-b)**. 41% of all GREs overlapped between the two datasets with thousands of GREs specific to either fetal or adult cortex **(Supplemental Figure 1c)**. To uncover the evolution of DNA sequences within these GREs, we identified all substitution events in the human lineage and human-ancestral lineages. Based on the availability of extant species genomes, we focused on 5 human-ancestral lineages: Human, Hominin, African great ape (A. G. Ape), Great ape (G. Ape) and Ape **(Figure 1a, Supplemental Figure 1d-h, Supplementary Table 1-2, Methods)**. As expected, we identified more substitutions on lineages with a longer branch length (i.e., longer evolutionary time) (Human, A. G. Ape, Ape) than lineages with a shorter branch length (Hominin and G. Ape) **(Figure 1b,d)**. Substitution rates were similar across the lineages after branch length normalization **(Figure 1c,e)**. To reveal cell type specific evolutionary patterns, we then counted the normalized substitutions within the GRE markers of major cell types **(Supplementary Table 3)**. To highlight differences across cell types per lineage, we divided values for each cell type to the mean value across all cell types (Methods). While there was a significant excess of substitutions in certain cell types compared to others for various lineages in the adult dataset, the fetal dataset revealed significantly higher substitutions only in microglia markers for all lineages **(Figure 1f-g)**. Together, these results reveal new patterns of cellular evolution within the human brain epigenome.

**Figure 1:**
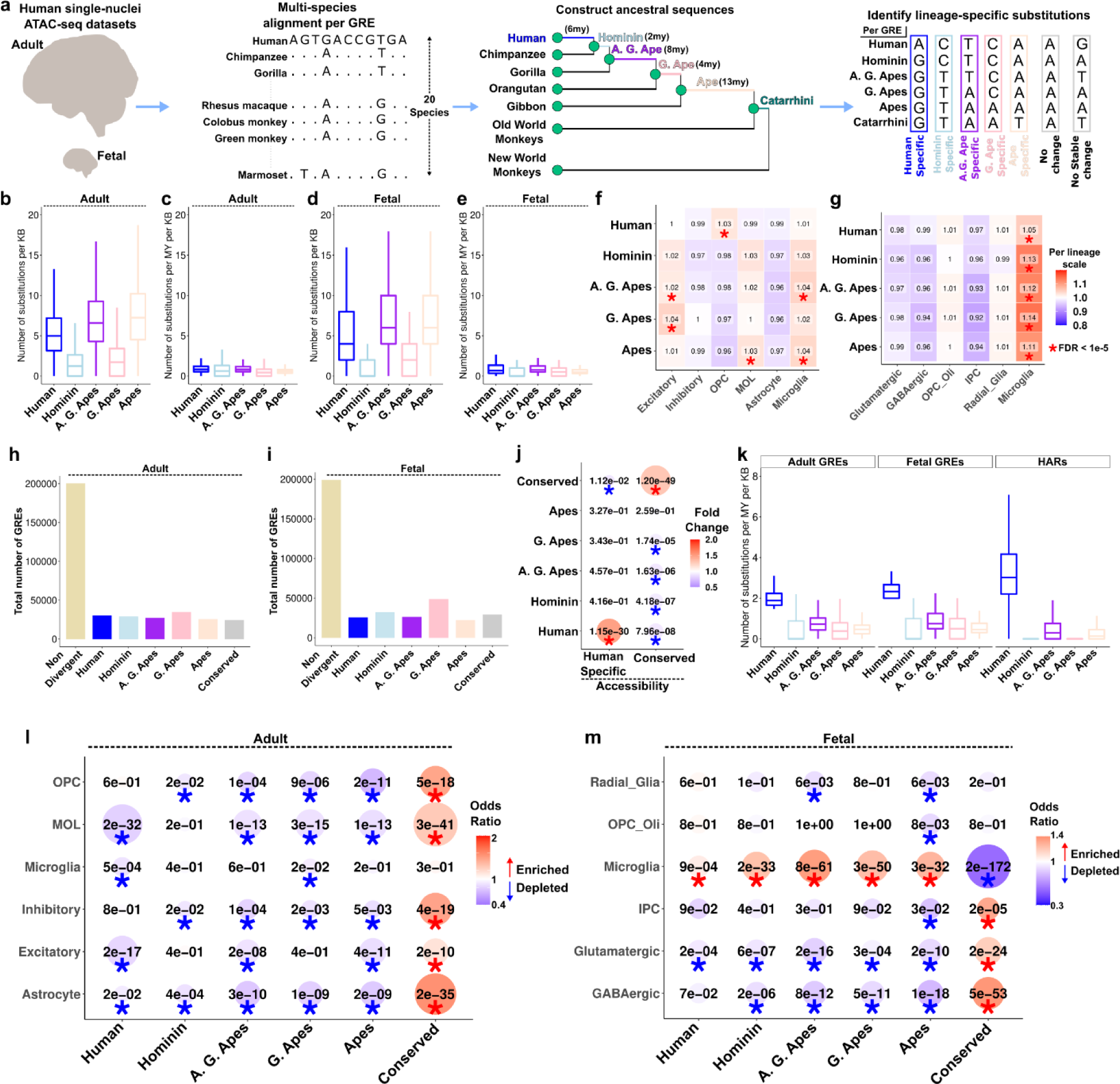
Identification of lineage-specific substitutions and divergent GREs. **(a)** Outline of the methodology to identify lineage-specific substitutions. **(b-e)** Substitution ratios within GREs per lineage. (**b,d**: GRE length normalized. **c,e**: GRE length and branch length normalized. **b,c**: adult, **d,e**: fetal). **(f-g)** Substitution ratios normalized to the mean value per lineage. Asterisks indicate FDR < 1e-5 (Chi-square test). **(h-i)** Number of GREs within each group in adult **(h)** and fetal **(i)** datasets. **(j)** Fisher’s exact test of overlaps between accessibility and substitution groups. Asterisk indicates FDR < 0.05. **(k)** Number of substitutions per million year per kb per lineage in adult, fetal and HAR datasets. **(l-m)** Fisher’s exact test of overlaps between cell type marker GREs and substitution groups. Asterisk indicates FDR < 0.05. Blue colors indicate depletions, red colors indicate enrichments.

### Identification of lineage divergent gene regulatory elements

Comparing relative substitutions across lineages, we classified ∼20,000-30,000 GREs that were either divergent in one lineage or conserved in all lineages, whereas most GREs were not classified as divergent or conserved **(Figure 1h-i, Supplementary Figure 1i-j, Supplementary Table 4, Methods)**. To test whether substitution-driven GRE classification is concordant with the functional measurements, we performed enrichments with species-specific accessibility changes in the adult dataset^16^. We found significant enrichments only between human-specific changes of accessibility and substitution or conserved accessibility and substitution **(Figure 1j)**. We compared lineage-specific substitutions in human-divergent GREs to HARs (shorter in length (∼250bp), fewer in number (∼3,000)^3,20^ and identified with a requirement of conservation across mammalian species^3^). We found that human-divergent GREs and HARs display a higher substitution rate in the human lineage and low substitution rate in ancestral lineages, although this difference is higher in HARs **(Figure 1k)**. Human-divergent GREs also overlap with both HARs and recently identified cortical HARs^16^ ∼10x more than ancestrally divergent GREs **(Supplementary Figure 1k)**. GC conversion ratio, typically high within accelerated regions^21^, is also higher in human-specific substitutions within HARs and cortical HARs than for all lineage-divergent GREs **(Supplementary Figure 1l)**. These results show that, lineage-divergent GREs show less divergence than accelerated regions but are more abundant to provide greater power for robust downstream analyses.

To complement our analysis of substitutions in cell type marker GREs, we asked whether cell type marker GREs are enriched in lineage-divergent GREs compared to the background of all GREs in the dataset. We note that in our previous analysis, we compared substitution rates across cell type marker GREs and did not test whether they were enriched compared to the background of all GREs (**Figure 1 f,g**). We found that cell type marker GREs are depleted in most lineage-divergent GREs and enriched in conserved GREs **(Figure 1l-m)**. Surprisingly, only fetal microglia marker GREs are uniquely enriched in lineage-divergent GREs, including humans, while depleted in conserved GREs, suggesting accelerated fetal microglia evolution in humans and their ancestors **(Figure 1l-m)**. We note that our approach overcomes the challenge of studying fetal brain tissue from non-human apes to reveal cellular evolution of the developing human brain.

### Identifying regulation of accelerated genes using multiomic datasets

To uncover genes that are regulated by lineage-divergent GREs, we sought to uncover the genes functionally linked to GREs. To achieve this in high throughput, we utilized adult and fetal cortical brain single-nuclei multiomic datasets and identified gene expression correlates of chromatin accessibility **(Supplementary Table 5)**. The GRE-gene linkage scores were significant for ∼40% of all GREs and were consistent across biological replicates in both datasets, indicating high reproducibility **(Supplementary Figure 2a-b)**. We then identified genes with significantly more divergent GREs, naming them regulation divergent genes (RDGs, Methods). We identified 30-130 RDGs per lineage in adult cortical brain, with limited overlap between lineages **(Figure 2a, Supplementary Figure 2c, Supplementary Table 6)**. Notable RDGs in both human and hominin lineages included *MSRA*, a methionine sulfoxide reductase implicated in both schizophrenia and autism^22^ and *CHD13* implicated in autism and attention deficit disorder^23^ **(Figure 2b)**. Moreover, human-divergent and hominin-divergent GREs linked to the same gene are distinct, indicating significant regulatory differences in both lineages **(Figure 2c-d)**. We also found a significant overlap with human-specific gene expression changes only for human RDGs but not for RDGs from ancestral lineages **(Figure 2e)**. We further identified RDGs in the fetal cortical brain that displayed low overlap with adult RDGs in all lineages **(Figure 2f-g, Supplementary Figure 2d-h, Supplementary Table 6)**. An example fetal-specific human RDG was *TFEB*, with 3 out of 8 of its linked GREs divergent in humans only in fetal tissue **(Figure 2h-i)**.

**Figure 2:**
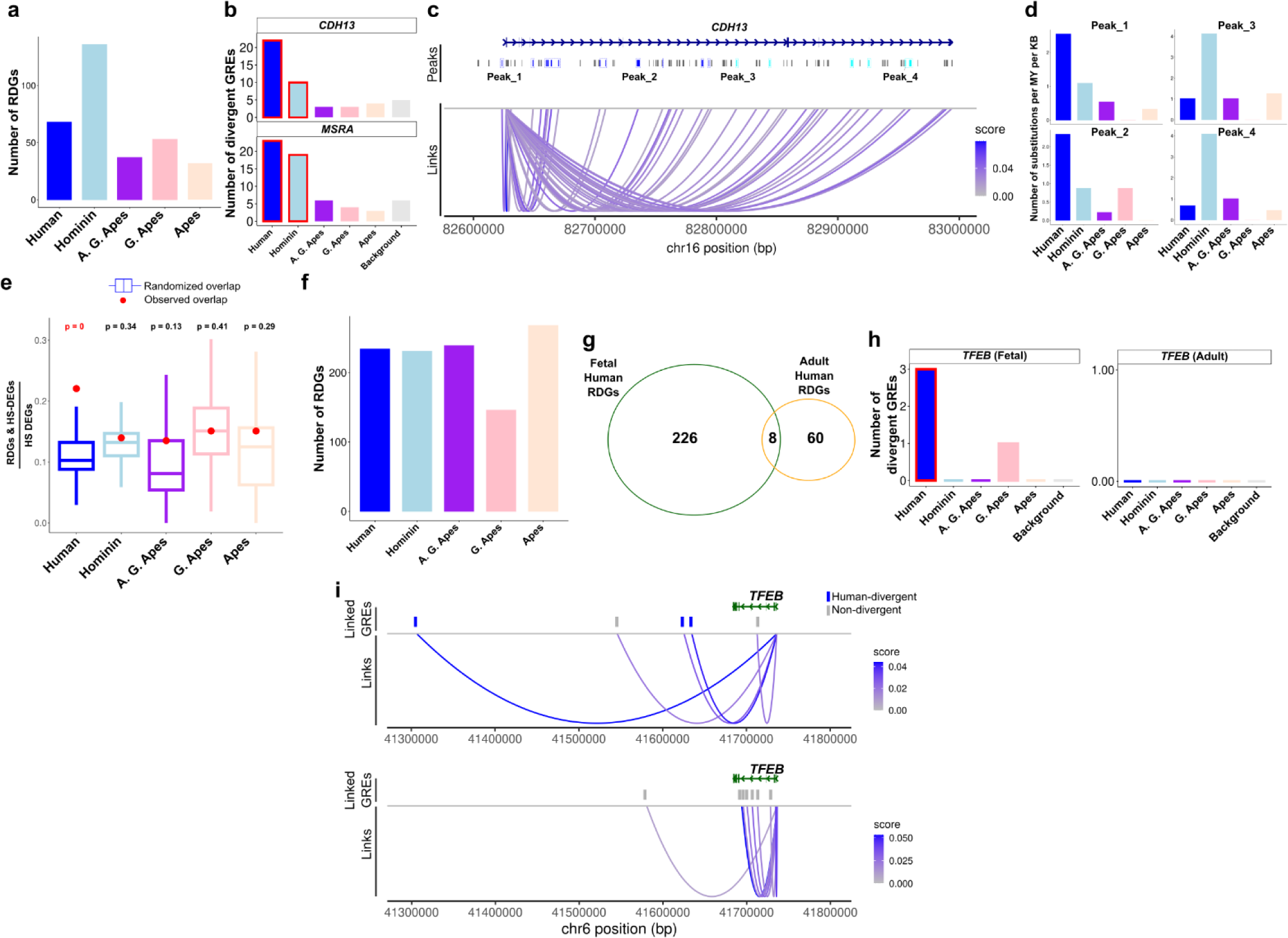
Identification of regulation divergent genes (RDGs). **(a)** Number of genes linked to significantly more divergent GREs compared to the background for each lineage. **(b)** Number of divergent GREs linked to *CDH13* and *MSRA* per lineage. Red rectangles indicate significant associations. **(c)** Coverage plot showing all GREs (labeled as Peaks) linked to *CDH13* expression. Blue colored peak regions indicate human-divergent GREs and light blue colored peak regions indicate hominin-divergent GREs. **(d)** Number of substitutions per million year per kb for the selected human-divergent and hominin-divergent GREs. **(e)** Empirical p-values of HS-DEG and RDG association per lineage (y-axis: ratio of HS-DEGs that are also RDGs among all HS-DEGs). Boxplots indicate randomized overlap (randomly selected ‘HS-DEGs’) repeated 1000 times. Red dot indicates the observed overlap. **(f)** Same as (a) but for fetal dataset. **(g)** Overlap between fetal and adult RDGs. **(h)** Number of divergent GREs linked to *TFEB* per lineage in fetal (left) and adult (right) datasets. **(i)** Same as **(c)** but for *TFEB* in fetal (top) and adult (bottom) datasets.

### Functional enrichments of RDGs and divergent-GREs

To investigate the fetal microglia identity divergence, we identified fetal RDGs linked to lineage-divergent microglia GRE markers and conducted gene ontology enrichments. This revealed 3 distinct enrichments: proteoglycan metabolic process, lymphocyte migration and intermediate filament-based process **(Figure 3a)**. While RDGs for proteoglycan metabolic process and intermediate filament-based process were contributed from all lineages, RDGs for lymphocyte migration were primarily from the human lineage **(Figure 3b-d)**. Proteoglycans are a major component in extracellular matrix that can affect neurite growth and myelination^24^ and microglia are known to be crucial for the homeostasis of extracellular matrix^24^. Notably, some human RDGs associated with the term lymphocyte migration are previously known to be markers of cytokine associated fetal microglia^25–27^ **(Supplementary Figure 3a-b)** and others (*ADAM8*, *DOCK8*, *PLEC*) are also associated with response to inflammation^28–30^, indicating human-specific regulation of cytokine-associated microglia in fetal brain development.

**Figure 3:**
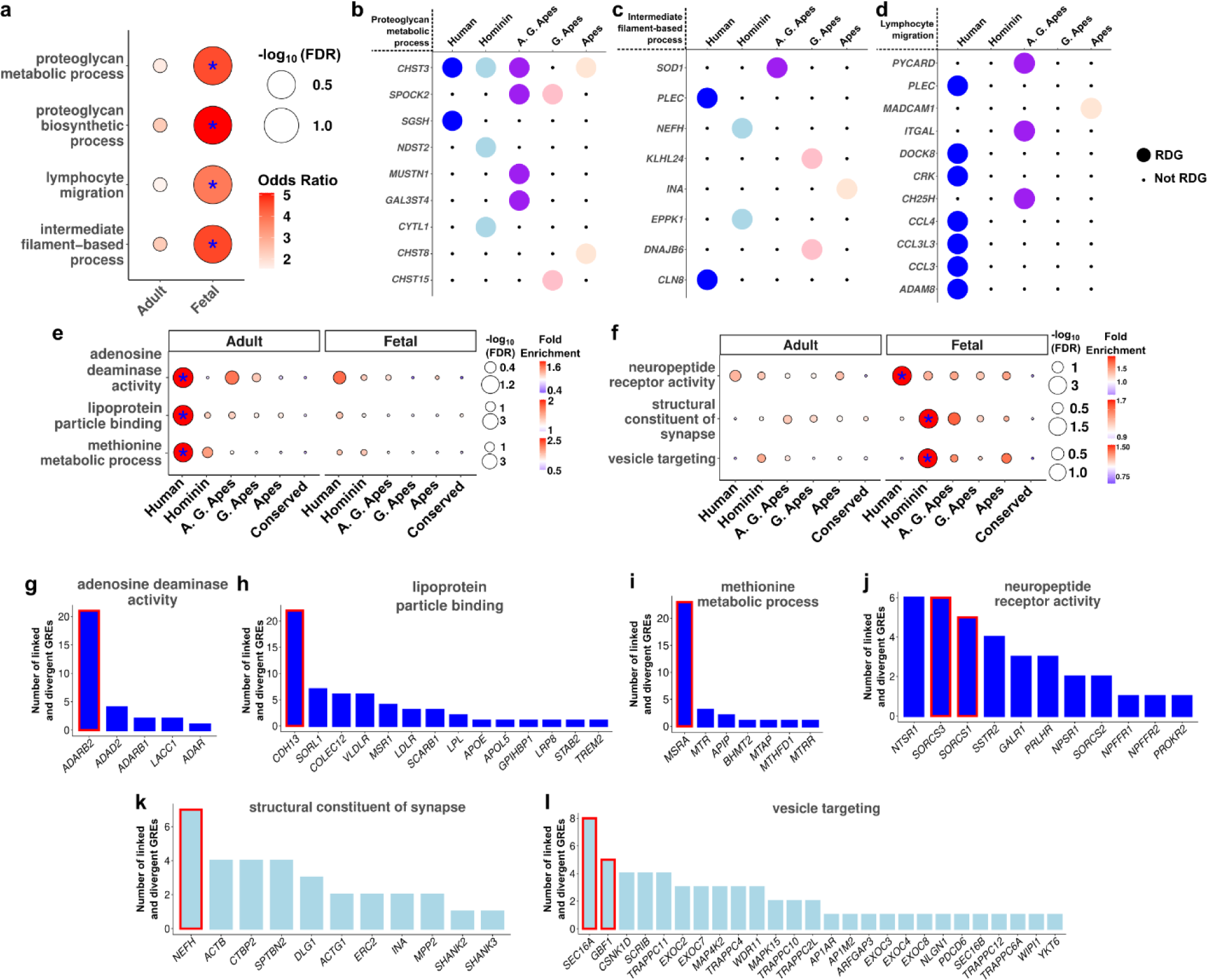
Functional enrichment analyses of divergent features. **(a)** Gene ontology enrichment results of RDGs linked to microglia marker divergent GREs. Asterisk indicates FDR < 0.05. **(b-d)** List of RDGs for each enrichment term per lineage. **(e-f)** Gene ontology enrichment results of GREAT for adult **(e)** and fetal **(f)** datasets. Each term is scaled separately with legends shown on the right side. Asterisk indicates FDR < 0.05 and fold enrichment > 1.3. **(g-l)** Number of divergent GREs for genes associated with the gene ontology term. Red rectangles indicate RDGs.

To identify functional enrichments of lineage-divergent GREs and RDGs, we performed gene ontology enrichments on lineage-divergent GREs (Methods). We found a small list of enrichments across most lineages in adult and fetal datasets **(Supplementary Table 7, Supplementary Figure 3c-d)**. After reducing the redundancy of gene ontology enrichment terms in the adult human lineage, we identified adenosine deaminase activity, lipoprotein particle binding and methionine metabolic process among the top enrichments highlighting metabolic pathways and RNA editing activity **(Figure 3e, Supplementary Figure 3c,e-f)**. In contrast, human and hominin lineage enrichments in the fetal datasets favored synapse structure and neuromodulation **(Figure 3f)**. To highlight the RDGs and other divergent GRE linked genes, we plotted genes and the number of divergent GREs linked to them for each of these gene ontology enrichments **(Figure 3g-l)**. Most terms were mainly driven by the GREs linked to RDGs although in all cases more than one gene (including non-RDGs) contributed to the overall enrichment. Our analysis implicates further biological pathways that may be lineage-specifically altered in the human brain evolution.

### Evolution of transcription factor binding sites in ape lineage

DNA sequence substitutions can alter transcription factor (TF) binding sites (TFBS)^31^. To provide a systematic investigation of the ancestral TFBS evolution patterns within human brain epigenome, we found gains and losses of all TFBS per lineage **(Supplementary Table 8, Methods)**. We created a binary matrix of motif presence per GRE across lineages and only considered stable changes as a TFBS gain or loss **(Figure 4a)**. We then reported the gain/loss ratio per TFBS as a readout to identify TFBS expansion/depletion across lineages **(Figure 4b)**. Total gain/loss ratios favored more gains than losses and were similar across lineages and datasets **(Supplementary Figure 4a-b**). Gain/loss ratios across TFBSs also showed high correlations between adult and fetal datasets for all lineages **(Supplementary Figure 4c)**. However, gain/loss ratio correlations across TFBSs revealed higher correlations between closer lineages, and lower correlations between more distant lineages **(Figure 4c-d)**.

**Figure 4:**
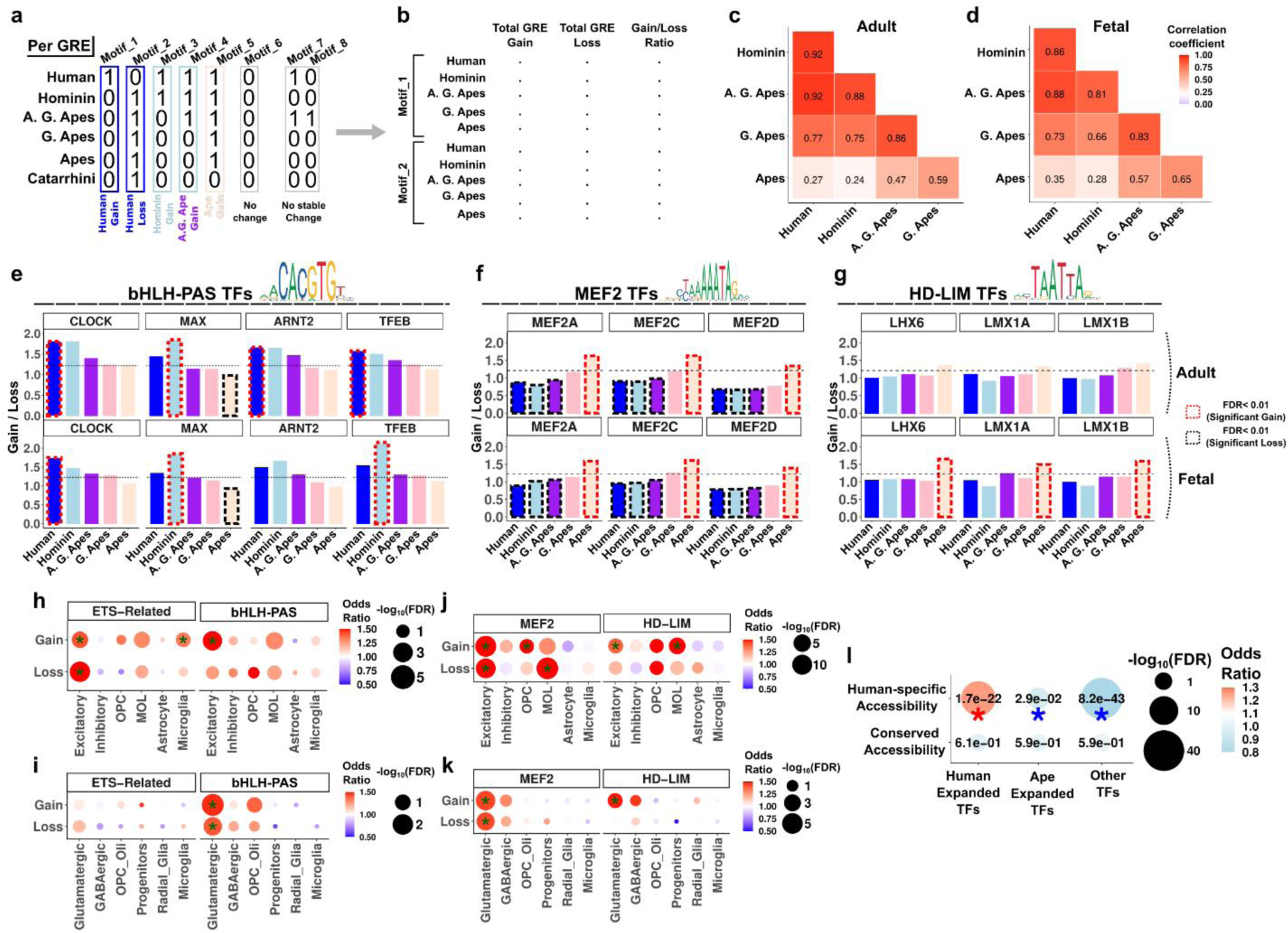
Transcription factor binding site evolution in human and ancestral lineages. **(a-b)** Identification of motif occurrences per lineage per GRE and total TFBS gains and losses per lineage. **(c-d)** Spearman rank correlations of gain / loss ratios across all motifs between lineages. **(e-g)** Gain / loss ratios of bHLH-PAS TFs **(e)**, MEF2 TFs **(f)** and HD-LIM TFs **(g)**. Dashed rectangles indicate significant expansions (red) and significant depletions (black). Horizontal dashed line indicates the global gain / loss ratio. **(h-k)** Fisher’s exact test (one-tailed) of gained or lost TFBSs in cell type marker GREs. Asterisk indicates FDR < 0.05. (l) Fisher’s exact test (two-tailed) of gained or lost (combined) TFBS changes in human-expanded TFs, ape-expanded TFs or other TFs with chromatin accessibility groups. Red asterisk indicates enrichment (FDR < 0.05), blue asterisk indicates depletion (FDR < 0.05).

We identified the significant TFBS expansions/depletions per lineage, by statistically assessing deviations from the background and other lineages **(Supplementary Table 9, Methods)**. Most expansions and depletions are detected in human, hominin and ape lineages **(Supplementary Figure 4d-g)**. Human-expanded TFBSs were similarly expanded in hominins and vice versa, although ape-expanded TFBSs displayed distinctly greater gain/loss ratio than all other lineages **(Supplementary Figure 4h, j).** We also detected a pattern of inverse correlation between human-hominin and ape lineages as expanded TFBSs in one group are relatively depleted in the other and vice versa **(Supplementary Figure 4h-k)**. Among the TF families with expanded TFBSs in human and / or hominin lineages, top enrichments were from bHLH-PAS factors, and ETS-related factors **(Figure 4e, Supplementary Figure 5)**. Notable TFs among them included CLOCK, a circadian clock gene with human brain evolution related functions^32^, MAX, a regulator of cell proliferation with unknown functions in the brain^33^, ARNT2, a regulator of activity dependent gene expression^34^ and TFEB; a regulator of oligodendrogenesis^35^. We additionally found that MEF2 factor TFBSs have expanded in ape lineage and depleted in human-hominin lineages whereas HD-LIM (homeodomain-LIM) factors have expanded in the ape lineage without significant depletion in other lineages **(Figure 4f-g)**. These findings reveal TFBS evolution patterns and provide a resource for all stable TFBS evolution events in the human brain epigenome **(Supplementary Table 8-9)**.

To reveal the cell type specificity of TFBS expansions, we then performed enrichments with cell type marker GREs and found that gains and losses are often enriched in the same cell type **(Figure 4h-k)**. However, there were also enrichments specific to expansions or depletions. For example, ETS-Related binding sites expanded in adult microglia and bHLH-PAS binding sites expanded in both adult and fetal glutamatergic cells **(Figure 4h-i)**. Among the ape expanded TFBSs, MEF2 binding sites were depleted in mature oligodendrocytes (MOL) and expanded in oligodendrocyte progenitor cells (OPCs) in the adult dataset, and HD-LIM binding sites were expanded in glutamatergic cells in both datasets as well as in mature oligodendrocytes **(Figure 4j-k)**. Considering that TFBS alterations affect chromatin accessibility^36^, we examined enrichments between GREs with altered TFBSs and human-specific chromatin accessibility changes. Targets of human expanded TFs were enriched in human-specific accessibility changes whereas targets of ape expanded TFs or other TFs were depleted **(Figure 4l)**. Taken together, these results identify TFs that significantly altered their target space in the evolution of human brain epigenome.

### Brain disease susceptibility in conserved and divergent GREs

Certain brain diseases such as schizophrenia, autism and Alzheimer’s are not reliably detectable in non-human primates, leading to the suggestion that human brain evolution contributed to the genetic susceptibility for these conditions^37–39^. While some studies statistically linked human-evolved genetic changes and disease susceptibility^40–45^, few contrasted this with conserved or non-human divergent features^41,43^. Interestingly, these studies favored greater enrichments of brain disease susceptibility in conserved genomic features compared to human-specific genomic features^41,43^. To provide a systematic comparison of disease susceptibility in human brain evolution, we performed LD score regression (LDSC) analysis^46^ for the conserved and divergent GREs as identified previously **(Figure 1, Supplementary Table 10)**. We performed regressions for the top 20,000 GREs to equalize the sample sizes (Methods). Strikingly, we found significant enrichments mainly among the conserved GREs in both adult and fetal brain **(Figure 5a)**. We reproduced this pattern with the top 10,000 or 5,000 GREs, indicating a robust trend **(Supplementary Figure 6a-d)**. Among the lineage-divergent enrichments, we mostly detected weak enrichments (0.01<FDR < 0.1) between the ancestral lineages and brain diseases / traits. We only observed two strong (FDR<0.01) enrichments between schizophrenia (SCZ) and hominin-divergent GREs and between intelligence (INT) and G. Ape-divergent GREs in the fetal brain **(Figure 5b)**. We did not identify any significant enrichments for the human divergent GREs **(Figure 5a-b)**. The total SNP numbers were similar across the groups, excluding the possibility of an excess number of SNPs driving the statistical differences **(Figure 5c-d)**. Sliding window analysis further showed that the enrichment comes from the GREs and not from the flanking regions included to obtain robust coefficient estimates^41,47^ **(Methods, Supplementary Figure 6e-f)**.

**Figure 5:**
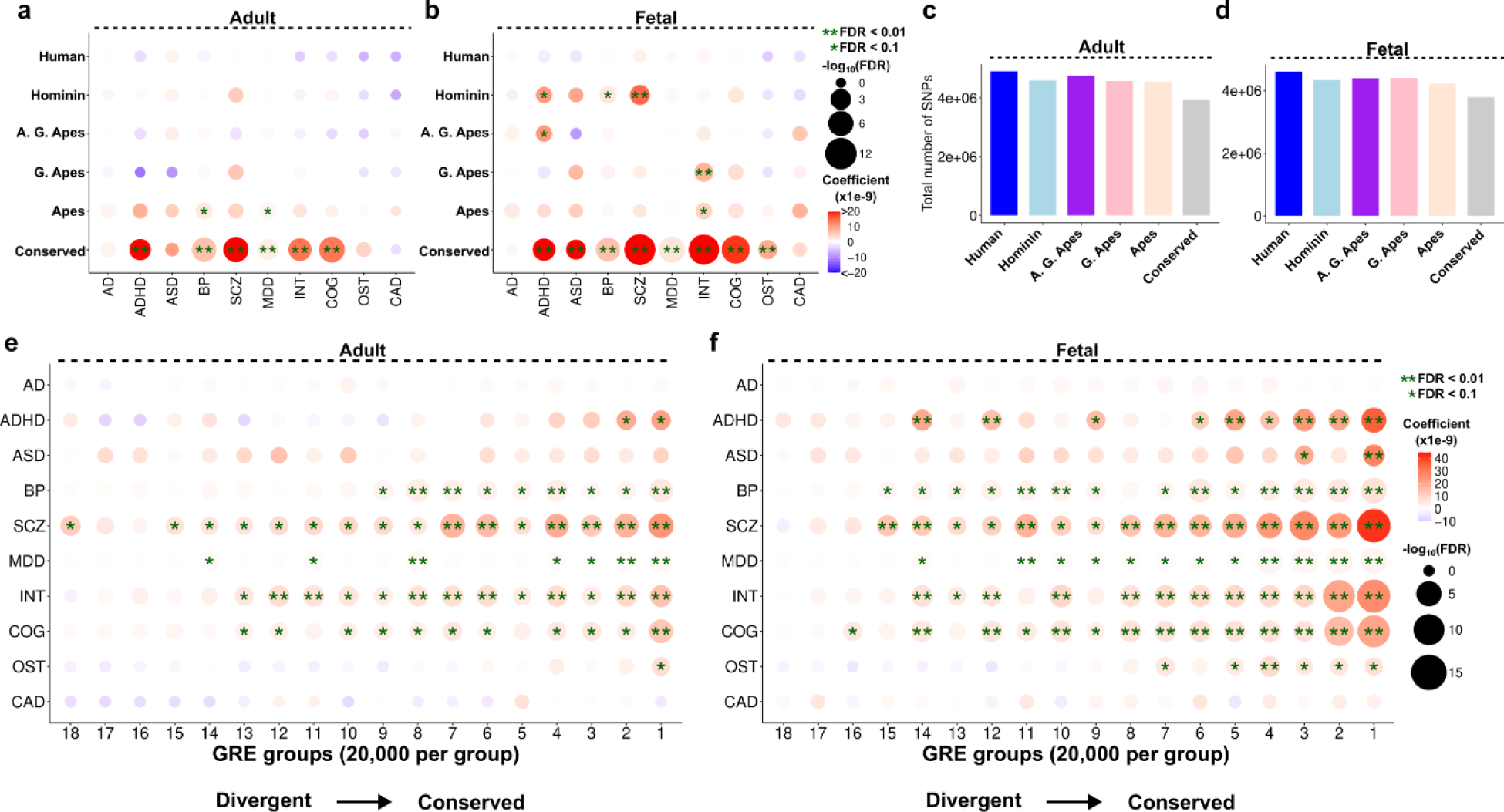
Association of disease variants and evolutionary divergence of GREs. **(a-b)** LDSC regression results between each disease category (x-axis) and lineage (y-axis) for adult **(a)** and fetal **(b)** datasets. Top 20,000 most divergent GREs were used to run the regressions in each lineage **(c-d)** Total number of SNPs mapping to the GREs per lineage per dataset (adult: **c**, fetal: **d**). **(e-f)** LDSC regression results for each GRE group (ordered based on conservation score and divided into 20,000 GREs per group) in adult **(e)** and fetal **(f)** datasets. In all panels, single asterisk indicates FDR < 0.1 and double asterisk indicates FDR < 0.01.

To test the association between conservation and disease susceptibility, we divided all GREs based on their average conservation score across 5 lineages, and similarly observed stronger enrichments among more conserved groups despite a similar total number of SNPs per group **(Figure 5e-f, Supplementary Figure 6g-h)**. Interestingly, fetal GREs often displayed more significant enrichments with greater effect size than conserved adult GREs **(Figure 5e-f)**. These results indicate greater disease susceptibility for more conserved GREs in human brain epigenome and motivate further discussion on the interplay between human brain disease susceptibility and human brain evolution.

## Discussion

We utilized DNA sequence substitutions within the cellularly resolved GREs to gain further insight into human brain evolution. Most cell type markers were enriched in conserved GREs except for fetal microglia markers **(Figure 1)**. Utilizing the multiomic datasets, we identified genes with significantly greater number of lineage-divergent GRE links (RDGs) **(Figure 2)** and used both divergent GREs and RDGs to elucidate further functional enrichments **(Figure 3)**. We also identified major TFBS evolution patterns across human and human-ancestral lineages, revealing lineage-specific expansions and depletions of all known TFBSs **(Figure 4)**. Finally, we found greater enrichments of brain disease susceptibility for the conserved GREs compared to divergent GREs **(Figure 5)**.

Fetal microglia markers were surprisingly more divergent than all other cell types **(Figure 1m)**. Interestingly cytokine and inflammatory response genes displayed regulatory divergence only among humans **(Figure 3d)**. Similarly, a previous study found increased expression of inflammatory genes in human microglia compared to rhesus macaque, marmoset and other species^48^. Other studies have established that neuroinflammatory genes are upregulated in aging, neurodegeneration, and development^25,26,49^. Our results suggest a human-specific divergence of cytokine-associated genes and inflammatory response genes in fetal microglia. However, fetal microglia GREs were also divergent in human-ancestral lineages. Recent studies on sequence divergence across numerous mammalian species also identified accelerated evolution in environmental response genes, especially immune responses^50,51^. Since microglial identity is driven by environmental stimuli^49^, fetal microglial may have particularly undergone accelerated evolution in humans and their ancestors.

Recent studies have identified species-specific gene expression changes at cell type resolution^15–17^. In this study, we identified genes linked to significantly more divergent GREs than expected by chance. Although RDGs show enrichment in human-specific gene expression changes, this approach complements transcriptomic assays, which offer snapshots of gene expression and may miss differential regulation by GREs. Single-nuclei transcriptomics can also mask an alternative start site usage due to being 3’ biased. An epigenome-driven approach is permissive to these mechanisms and augments transcriptional data. In summary, RDGs offer an in-silico alternative to transcriptomic comparisons, expanding our ability to detect species-specific gene regulation.

Analysis of thousands of TFBS evolution can reveal lineage-specific patterns and offer unique insights about regulatory evolution^50,52^. Despite tissue and cell type specificity of TFBSs, TFBS evolution in human brain is poorly understood. In this study, we systematically identified gain and loss events and observed that correlations of TFBSs gain/loss ratios roughly recapitulates phylogeny **(Figure 4c-d)**. Analysis of TFs with expanded or depleted TFBS pools primarily revealed significant changes in human-hominin and ape lineages, suggesting widespread expansion or contraction of certain TF targets. Since TFs regulate critical developmental decisions, TFBS expansions might be tied to the evolution of cell type identity. Indeed, many of the newly gained TFBSs were significantly associated with cell type markers while their newly lost targets were not **(Figure 4h-k)**. We provide the list of TFBS gains and losses in human and human-ancestral lineages **(Supplementary Table 8-9)**.

Evolutionarily conserved molecular features are more associated with disease risk than evolutionarily divergent molecular features^53,54^. However, diseases can be species-specific, indicating that the adaptive benefit of newly evolved features must exceed the deleterious effects they might have entailed^55,56^. Some human brain diseases are also thought to be linked to human brain evolution, although disease association in human-divergent features are rarely contrasted with conserved features or non-human divergent features^41,43^. To our knowledge, no previous study systematically compared the human lineage to its ancestral lineages for disease susceptibility and contrasted this with the conserved features. We investigated this specifically for the adult and fetal cortical brain GREs and uncovered brain disease susceptibility enrichments mostly among conserved features, which increased with greater conservation score **(Figure 5)**. Strikingly, human divergent-GREs were not enriched in disease susceptibility, and we detected enrichments only for human-ancestral divergent GREs **(Figure 5)**. Therefore, we could not find a human lineage-specific susceptibility to the brain diseases we examined. We note, however, that the disease variants in our study are mainly comprised of common variants identified through genome-wide association studies that do not have enough statistical power to capture the rare variants that can target distinct genomic regions^57^. It is therefore possible that rare disease variants might have greater association with more divergent GREs. Alternatively, human evolution may have rendered the human brain more susceptible to certain diseases not by disruption of the same biological pathways but by increasing the human brain physiology vulnerability to certain disruptions secondarily through other major evolutionary changes.

## Acknowledgments

The authors thank Dr. Soojin V. Yi for their critical comments on the manuscript. G.K. is a Jon Heighten Scholar in Autism Research and Townsend Distinguished Chair in Research on Autism Spectrum Disorders at UT Southwestern. E.C. is a Neural Scientist Training Program Fellow in the Peter O’Donnell Brain Institute at UT Southwestern. This work was partially supported by the James S. McDonnell Foundation 21^st^ Century Science Initiative in Understanding Human Cognition Scholar Award (220020467), the Simons Foundation (947591), NHGRI (HG011641), NINDS (NS115821, NS126143) and NIMH (MH126481, MH103517) to G.K.

## Author Contributions

E.C. conceptualized the study, performed all analyses and wrote the manuscript. G.K. provided guidance and supervision. E.C. and G.K. edited the manuscript.

## Data and Code Availability

All datasets utilized in this study are publicly available with the following accession numbers: adult single-nuclei ATAC-seq: GSE192774, fetal single-nuclei ATAC-seq: GSE162170, adult single nuclei multiome (ATAC+RNA): GSE207334, fetal single-nuclei multiome (ATAC+RNA): GSE162170, UCSC multi-species alignment: https://hgdownload.soe.ucsc.edu/goldenPath/hg38/multiz30way/

## Methods

### Identification of lineage-specific substitutions

Since we are interested in the substitutions specific to ape lineage and its derived lineages extending to the human lineage, we focused on ape, old world monkey and new world monkey species for our comparisons. We therefore extracted the following genomes from UCSC 30-way alignment dataset^58^: Ape species: *hg38, panTro5, gorGor5, ponAbe2,* nomLeu3. Old world monkey species: rheMac8*, macFas5, macNem1, cerAty1, papAnu3, chlSab2, manLeu1, nasLar1, colAng1, rhiRox1, rhiBie1.* New world monkey species: *calJac3, saiBol1, cebCap1, aotNan1.* We also only retained these species within the associated phylogenetic tree. We then extracted each GRE from this multi-species alignment by removing all sequences absent in the human genome, essentially removing all potential human-specific deletions (*strip.gaps.msa* from *rphast*^59^). Per GRE, we then computed the maximum likelihood for the tree with *pml* function and optimized it for F81 substitution model using *optim.pml* function using *phangorn*^60^. Then we estimated the ancestral sequence probabilities using maximum likelihood approach with *ancestral.pml* function. We only considered ancestral states estimated with a probability > 0.75 for one of the nucleotides. If no nucleotide exceeded a probability of 0.75, we assigned the ancestral state as N (instead of A, C, T or G). These positions were discarded during the identification of the lineage-specific substitutions (see below). With this approach, we could reconstruct ancestral sequences within apes, as well as the ancestral state of the entire ape lineage since we have old world monkeys as its sister group and new world monkeys as the outgroup. Since we are only focused on the ancestral sequences within apes and how they compare with human sequence, we proceeded with the following sequences: human, hominin-ancestral, African great ape-ancestral, great ape-ancestral and ape-ancestral.

We then identified substitutions that occurred in each lineage with the following criteria:

1. Ancestral sequence should be confidently reconstructed in all lineages for consideration for a substitution (i.e., any sequence tagged with ‘N’ was not considered).
2. Substitutions that occurred more than once across the 5 lineages were discarded (e.g., Human: A, Hominin: **C**, African Great Ape: **C**, Great Ape: **G**, Ape: **G**).
3. Similarly, substitutions that were reversed in a daughter lineage were also discarded (e.g., Human: **C**, Hominin: **C**, African Great Ape: **A**, Great Ape: **C**, Ape: **C**).
4. We then only retained the substitutions that putatively occurred once across the 5 lineages (e.g., Human: **T**, Hominin: **T**, African Great Ape: **T**, Great Ape: **A**, Ape: **A**).

We created a list of all substitutions across the 5 lineages including which ancestral node the substitution took place, the ancestral nucleotide, the derived nucleotide, the position of the nucleotide in the human genome (hg38), the GRE that contains the substitution and the position of the substitution within the GRE **(Supplementary Tables 1-2)**.

To calculate the total number of substitutions, we summed all substitutions per lineage per GRE. To normalize for the branch length of each lineage (i.e., evolutionary time), we divided these sums to their corresponding branch length in million years (lower-bounds of a previous phylogenetic estimate^61^; Human: 6, Hominin: 2, A.G. Ape: 8, G. Ape: 4, Ape: 13). To further adjust for the length of the GRE and calculate substitution rate per kilobase, we then divided these values to the length of the GRE (in base pairs) and multiplied with 1000. This yielded normalized substitution rate per million years per kilobase (per MY per kb) for each GRE for human lineage and human-ancestral lineages.

We note that we performed our analysis on an adult and a fetal cortical brain dataset. While these datasets do not have resolution of all cortical regions (adult: posterior cingulate cortex, fetal: cerebral cortex), our previous analyses showed that – at least in the adult brain –, there is a very high (∼90%) overlap of GREs between datasets from different cortical regions^16^ indicating high reproducibility of DNA-based substitution results across cortical brain datasets by definition. We have therefore prioritized our dataset selection based on cellular resolution, multi-species comparison, and multiomic dataset availability with similar age and from a similar tissue^16,18,19^.

### Cell type marker GREs

We grouped nuclei into broad cell type categories in both fetal and adult datasets. We then aggregated counts per sample for each of the broad cell type category. To identify GREs that are significantly more accessible (i.e., cell type marker GREs) per cell type, we performed differential analysis using edgeR on the aggregated matrices. GREs that were more accessible for the given cell type compared to other cell types were identified with FDR < 0.05 and logFC > 0.3 cutoffs **(Supplementary Table 3)**. This analysis was performed separately for the fetal and adult datasets for the cell type categories described in **Supplementary Figure 1a-b**.

### Cell type marker analysis of the substitutions

To understand the substitution patterns within the cell type markers, we summed evolutionary time normalized substitution values for all marker GREs per lineage per cell type. These values were then normalized for length by dividing them to the total length of all marker GREs per cell type. To highlight deviations of normalized substitutions across cell types, we then divided each value to the mean value across all cell types for each lineage. This yielded values around 1 with values >1 enriched in substitutions compared to other cell types for each lineage. We tested the statistical significance of enrichments with a one-sided Chi-square test. P-values were adjusted for multiple testing using FDR. Results with FDR < 1e-5 were considered a significant deviation.

### Evolutionary classification of the GREs

To classify GREs based on their substitution differences across lineages, we implemented three different cutoffs. First, we randomly sampled 10,000 GREs 1,000 times and calculated a background proportion of substitutions across lineages for each randomly selected 10,000 GREs. This yielded 1,000 proportions across lineages. We then calculated the proportion for each GRE separately and empirically calculated the p-value as the number of times the background proportion exceeded the observation for each lineage divided to 1,000 (subsequently adjusted for multiple testing using FDR). We identified the fold change by dividing the observed proportion to the median value of the randomly sampled proportions. We identified GREs divergent in a given lineage with cutoffs of FDR < 0.05 and fold change > 1.5. Second, we z-transformed the substitution proportions of all GREs per lineage and only retained significant GREs if their standard deviation is >1 for the lineage in which they are significantly more divergent. This further highlighted GREs that contain a substantial number of substitutions for the given lineage compared to other GREs. Third, we further filtered all divergent GREs to contain at least 2 substitutions for the lineage in which they are divergent. We did not require a GRE to be divergent in only one lineage with these filters. However, we still detected very low Jaccard similarity index (<0.05) of lineage-divergent GREs between any two lineages indicating very low to no overlap of divergent-GREs between lineages.

To identify conserved GREs, we found all GREs with normalized substitution value lower than at least 50% of the GREs for each lineage. We then retained the intersection of these GREs among all lineages. We additionally required this list to not contain any previously identified lineage-divergent GREs. We named the resulting list ‘conserved GREs’.

### Enrichment with species-specific accessibility changes

We combined the human-specific chromatin accessibility changes (compared to chimpanzee and rhesus macaque) across all cell types from a recent comparative single-nuclei ATAC-seq study (human dataset of this reference is also the adult dataset we used in this study)^16^. We defined conserved chromatin accessibilities as GREs that did not display a species-specific (across human, chimpanzee, rhesus macaque) accessibility in any cell type. To perform enrichment while accounting for GRE length, we performed a logistic regression with accessibility classification as the response variable (Human-specific or others. Conserved or others), and GRE length and substitution classification as the predictor variables (Human-divergent or others, Hominin-divergent or others etc.). P-value was computed with Wald’s test and FDR was calculated for multiple testing correction. Results with FDR < 0.05 were considered significant enrichments. We note that this enrichment was only done with the adult dataset since comparative single-nuclei accessibility results are from adult cortical brain tissues.

### Comparisons with HARs

Previously published HARs^42,62–65^ and cortical HARs^16^ were overlapped with lineage-divergent GREs using bedtools^66^. We identified lineage-specific substitutions within these regions as described above. To compute the GC conversion ratio, we divided the total number of conversions from A/T to G/C to all detected substitutions per HAR and lineage-divergent GRE.

### Cell type marker enrichment of lineage-divergent GREs

We performed two-tailed Fisher’s exact tests to determine whether overlaps between cell type marker GREs and lineage-divergent / conserved GREs are significantly enriched / depleted. We used all GREs for each dataset (adult or fetal) as the background. Overlaps with FDR < 0.05 and odds ratio > 1 were labeled as enriched and overlaps with FDR < 0.05 and odds ratio < 1were labeled as depleted.

### Identification of regulation accelerated genes

To identify GRE – gene expression linkage, we utilized multiome datasets obtained from fetal and adult brains^18,19^. Since the adult brain multiome dataset is from a different cortical brain region (prefrontal cortex instead of posterior cingulate cortex in the cross-species dataset), we re-generated the multiome GRE-cell count matrix on our set of GREs using *FeatureMatrix* from *Signac*^67^. Unsurprisingly, total read counts were tightly correlated between the original and the newly generated matrices (Spearman’s rho ∼= 0.99) since there is a high degree of overlap between GREs from different adult cortical regions. We then calculated potential GRE – gene expression links using *LinkPeaks* function from Signac that utilizes correlation between accessibility and gene expression of nearby genes and compared with the randomly selected associations^67,68^. Significant links were identified with FDR < 0.05 and score > 0.01 cutoffs. We performed this analysis separately for fetal and adult datasets.

To identify genes that are linked to a significant number of more lineage-divergent GREs, we first randomly selected the same number of GREs for each lineage-divergent GRE group (e.g., Human-divergent GREs) and found the number of linked GREs for each gene. This was repeated 1000 times to create a background. We then performed the same operation for lineage-divergent GREs (e.g., Human-divergent GREs) and identified the genes with significantly more linked lineage-divergent GREs than the background (FDR < 0.05). We additionally required these genes to be linked to at least 2 lineage-divergent GREs and at least 5 total GREs to increase the robustness of the results. We performed this analysis separately for each lineage. The final list of genes with significantly more divergent GRE links were referred to as regulation accelerated genes (RDGs). We performed this analysis separately for fetal and adult datasets.

To test the overlap between RDGs and human-specific differentially expressed genes (i.e., HS-DEGs), we combined HS-DEGs (from ref.^16^) across all cell types and created a unique list of HS-DEGs. To determine whether the overlap between RDGs and HS-DEGs is significant, we created a background of randomly selected ‘HS-DEGs’ and tested whether the observed overlap is significantly (p<0.05) greater than the background overlap. This analysis was repeated for RDGs from each lineage.

### Gene ontology enrichments

For gene ontology enrichment on divergent microglia markers, we found all microglia marker GREs that are divergent in at least one lineage. We then found RDGs linked to these GREs. To highlight functional enrichments within microglial identity, we used all genes linked to all microglia marker GREs as the background. Gene ontology enrichment was performed with *clusterProfiler*^69^ and terms with adjusted p-value < 0.05 were considered as significant enrichments. We performed this enrichment separately for fetal and adult datasets.

To perform gene ontology enrichments on the genomic regions, we aimed to capture enrichments that are associated with both divergent GREs and RDGs. To achieve this, we only retained GREs that are linked to at least one gene. Additionally, to highlight enrichments that differ across the lineages, we set the background as all lineage-divergent GREs (plus conserved GREs to provide an additional contrast) and foreground as GREs divergent in one lineage. We then performed gene ontology enrichment with rGREAT^70,71^ and considered terms with adjusted p-value < 0.05 and fold enrichment > 1.3 significant. We additionally required each enriched term to be associated with at least 10 divergent GREs and at least 1 RDG for the given lineage for increased robustness of the results. Both requirements filtered an additional >50% of the enrichment terms in each lineage. We then reduced the redundancy of the final list of enriched terms using *rrvigo* (Revigo package in R language)^72,73^.

### Identification of TFBS evolution patterns across lineages

To identify motif occurrences in the human and human-ancestral sequences, we downloaded the JASPAR 2022 non-redundant *H. sapiens* motif dataset and found motif occurrences in all GREs for each lineage using *matchMotifs* and *motifMatches* functions from *motifmatchr* package^74^. This yielded a binary matrix of motif presence / absence per lineage and per motif. We performed this analysis for all GREs. For each motif, we scanned GREs to find the lineages that gained / lost the motif. Similar to substitutions, we only considered the gain/loss of motifs if they were gained / loss exactly once. Gain / loss was assigned to the lineage the change occurred. To understand the putative TFBS expansion / depletion across the entire epigenome, we counted gains and losses per motif across all GREs and divided total gains to total losses per lineage.

Significant expansion / depletion of a TFBS across lineages was identified by Chi-square comparison of gain / loss ratio to i) overall gain / loss ratio of all TFBS in all lineages and ii) gain / loss ratio in other lineages for the given TFBS. To consider deviations significant, FDR < 0.01 was required for both comparisons in the same direction (>1 for expansions, <1 for depletions). We further filtered the TFBS for the accessibility / expression of their TF in at least 25% of the cells in at least one cell type category using the original annotations from the reference datasets^16,19^.

Enrichment with accessibility changes were performed as described above for lineage-divergent GREs except with putatively gained + lost targets of human-expanded TFs, ape-expanded TFs or other TFs.

### Cell type enrichment of TFBS evolution patterns

GREs that gained / lost a TFBS for the given TFBS were tested for enrichment in cell type marker GREs using two-tailed Fisher’s exact test. Enrichments were determined with FDR < 0.05 and odds ratio > 1.3 cutoffs. Since many TFBSs are from the same family and with similar motifs to each other, we also combined all TFBS gains / loss for some of these families (MEF2, bHLH-PAS, ETS-related, HD-LIM) **(Supplemental Figure 5)** and performed the same analysis to elucidate more robust cell type evolution patterns **(Figure 4h-k)**.

### LDSC regression

To perform LD (linkage-disequilibrium) score (LDSC) regression analysis, we expanded each GRE 25kb both upstream and downstream. We ranked GREs based on divergence using fold change of divergence across lineages. Conservation score was calculated as 1 / mean fold change across all lineages per GRE. Top 20000, 10000 or 5000 GREs were selected to perform LDSC regressions.

Genome-wide association study (GWAS) summary statistics from brain disorders (ADHD^75^: attention-deficit/hyperactivity disorder, ASD^76^: autism spectrum disorders, BP^77^: bipolar disorder, SCZ^78^: schizophrenia, MDD^79^: major depressive disorder, AD^80^: Alzheimer’s disease), brain traits (INT^81^: intelligence, COG^82^: cognitive function) and non-brain disorders (OST^83^: osteoporosis, CAD^84^: coronary artery disease) were downloaded and arranged for use in LDSC regression using *munge_sumstats*^46^. LDSC regressions were then performed with the recommended parameters^46^. All GREs (separately for fetal and adult datasets) were used as a background in all regressions. Results with FDR < 0.1 were considered weak enrichments and results with FDR < 0.01 were considered strong enrichments.

To assess the contribution of 25kb flanking regions in LDSC regression statistics, we performed the same analysis by shifting the 50kb window in 5kb increments from −100kb to +100kb relative to the center of the GRE, similar to a previous study^41^.

---

**Supplementary Figure 1:**
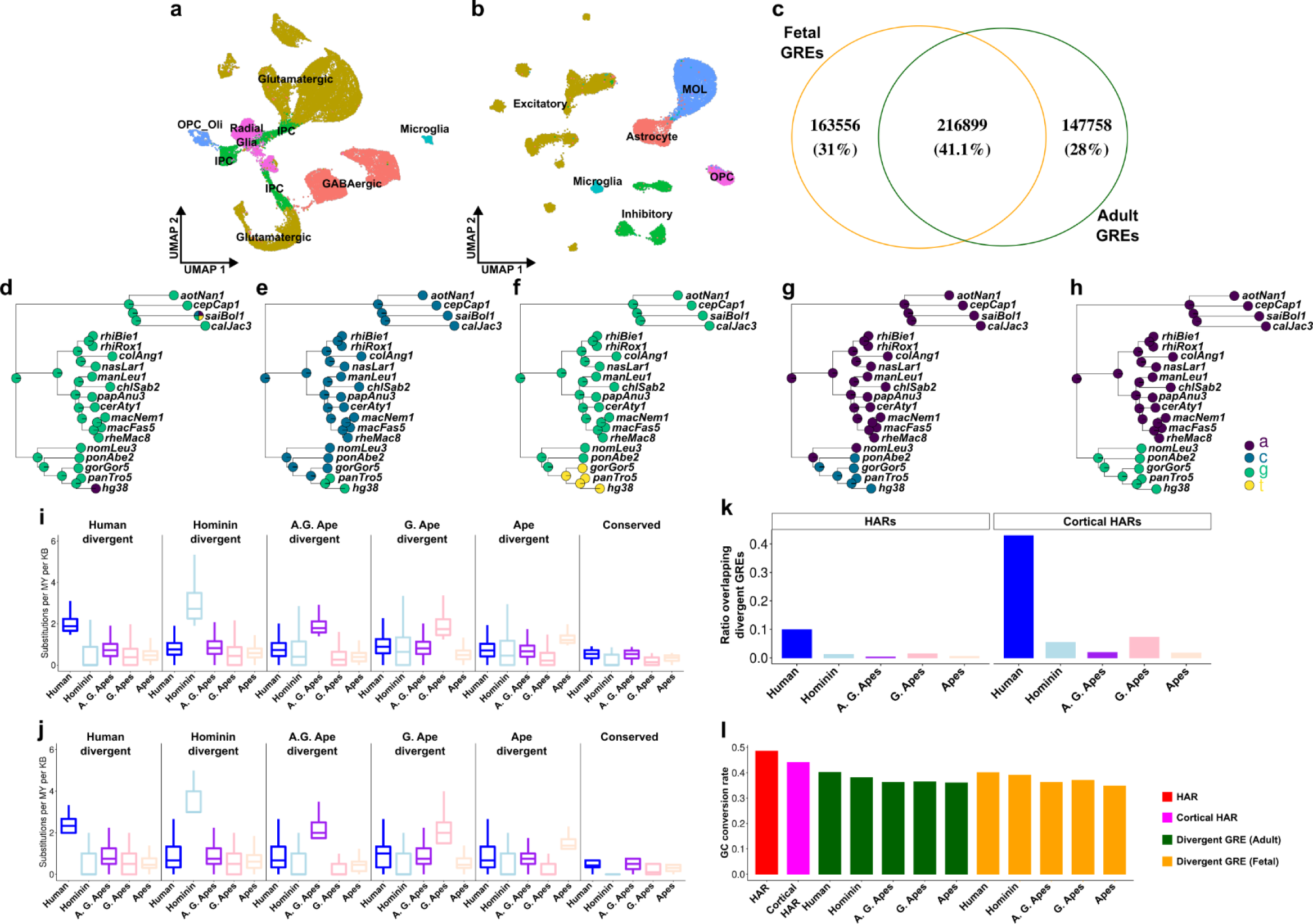
Additional plots on the datasets, substitutions and divergent GREs. **(a-b)** UMAP plots of fetal **(a)** and adult **(b)** datasets. **(c)** Overlap of GREs between adult and fetal datasets. **(d-h)** Phylogenies depicting lineage-specific substitutions in human **(d)**, hominin **(e)**, A.G. Ape **(f)**, G. Ape **(g)** and ape **(h)** lineages. Circles correspond to species included in the dataset and ancestral nodes with a reconstructed sequence. **(i-j)** Boxplots of substitutions per million year per kb for all lineage-divergent GREs and conserved GREs. **(k)** Ratio of HARs / cortical HARs overlapping divergent GREs. **(l)** Ratio of GC conversion in HARs, cortical HARs and lineage-divergent GREs in adult and fetal datasets.

**Supplementary Figure 2:**
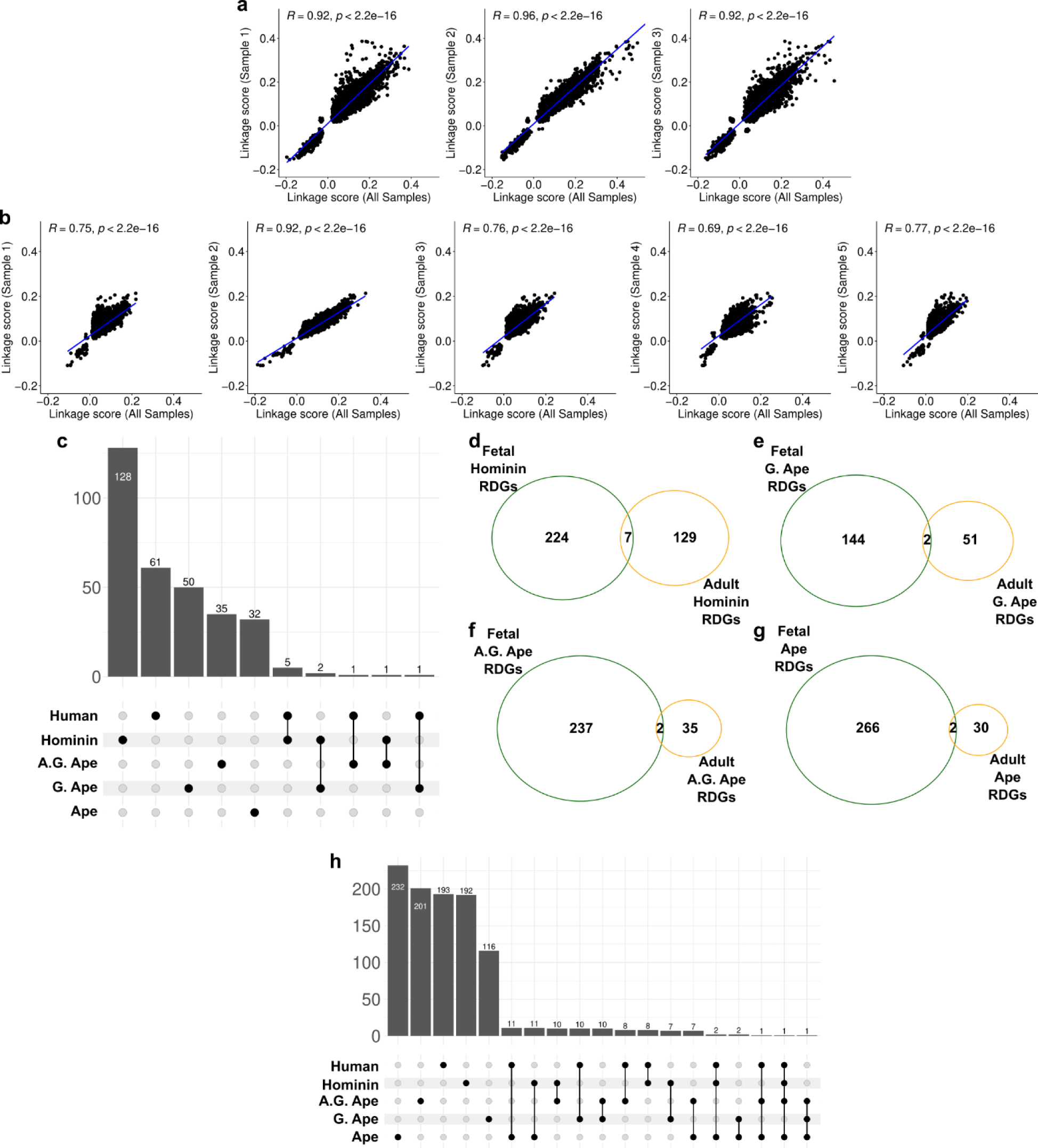
Supporting analyses on regulation divergent genes. **(a-b)** Spearman rank correlations of gene linkage scores between biological replicates in fetal **(a)** and in adult **(b)** datasets. **(c)** Number and overlap of RDGs across lineages in adult dataset. **(d-g)** Overlap of RDGs between fetal and adult datasets per lineage. **(h)** Same as **(c)** but in fetal dataset.

**Supplementary Figure 3:**
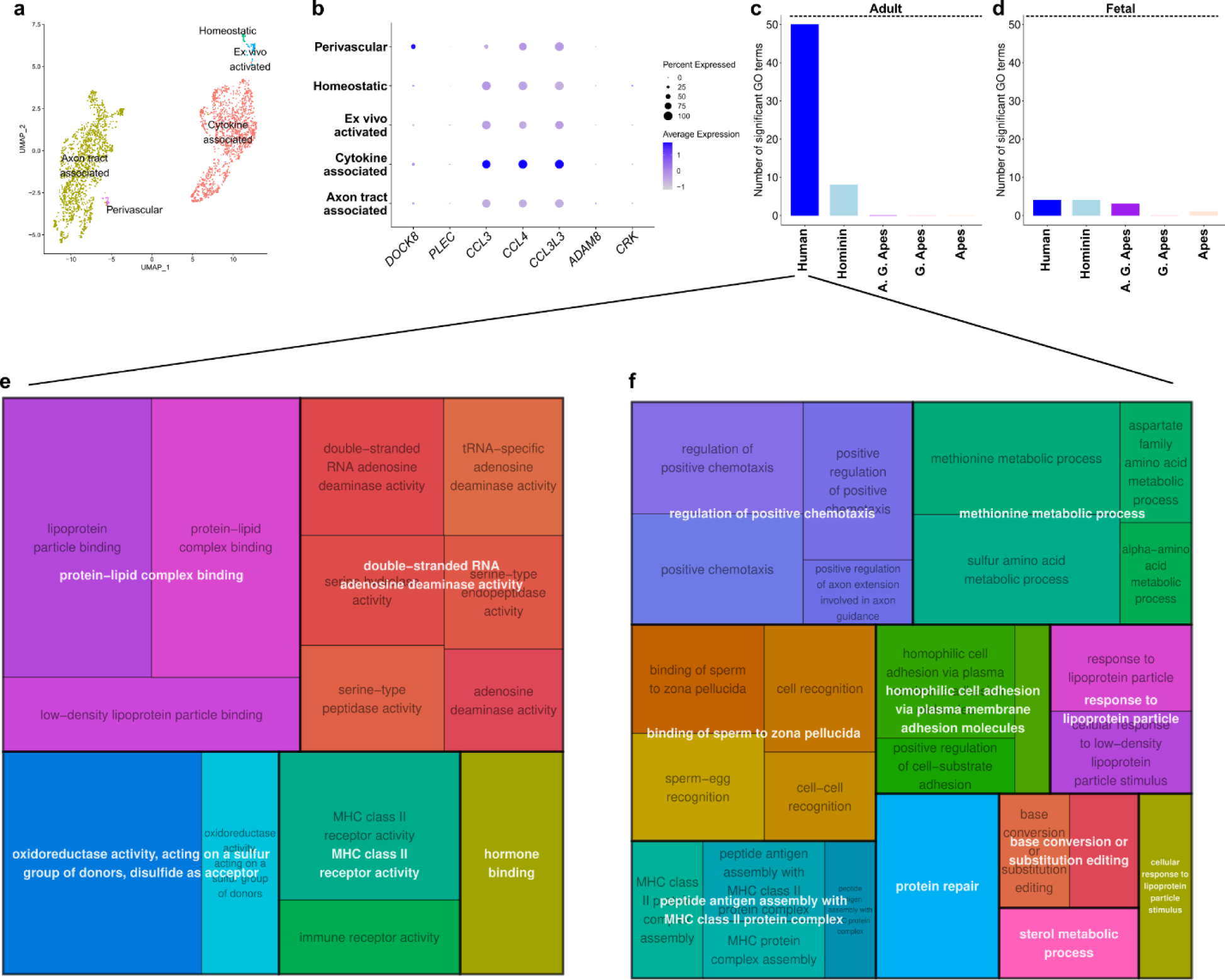
(a) UMAP of annotated fetal microglial cells. **(b)** Normalized expression values of human RDGs across microglial cell types. Note that while some genes have low expression levels, these RDGs are linked to at least one microglia marker GRE and might alter expression status. **(c-d)** Number of significant GO terms associated with divergent features within each lineage. **(e-f)** Revigo plots that reduce the redundancy of human associated enrichments (molecular function **(e)** and biological process **(f)**).

**Supplementary Figure 4:**
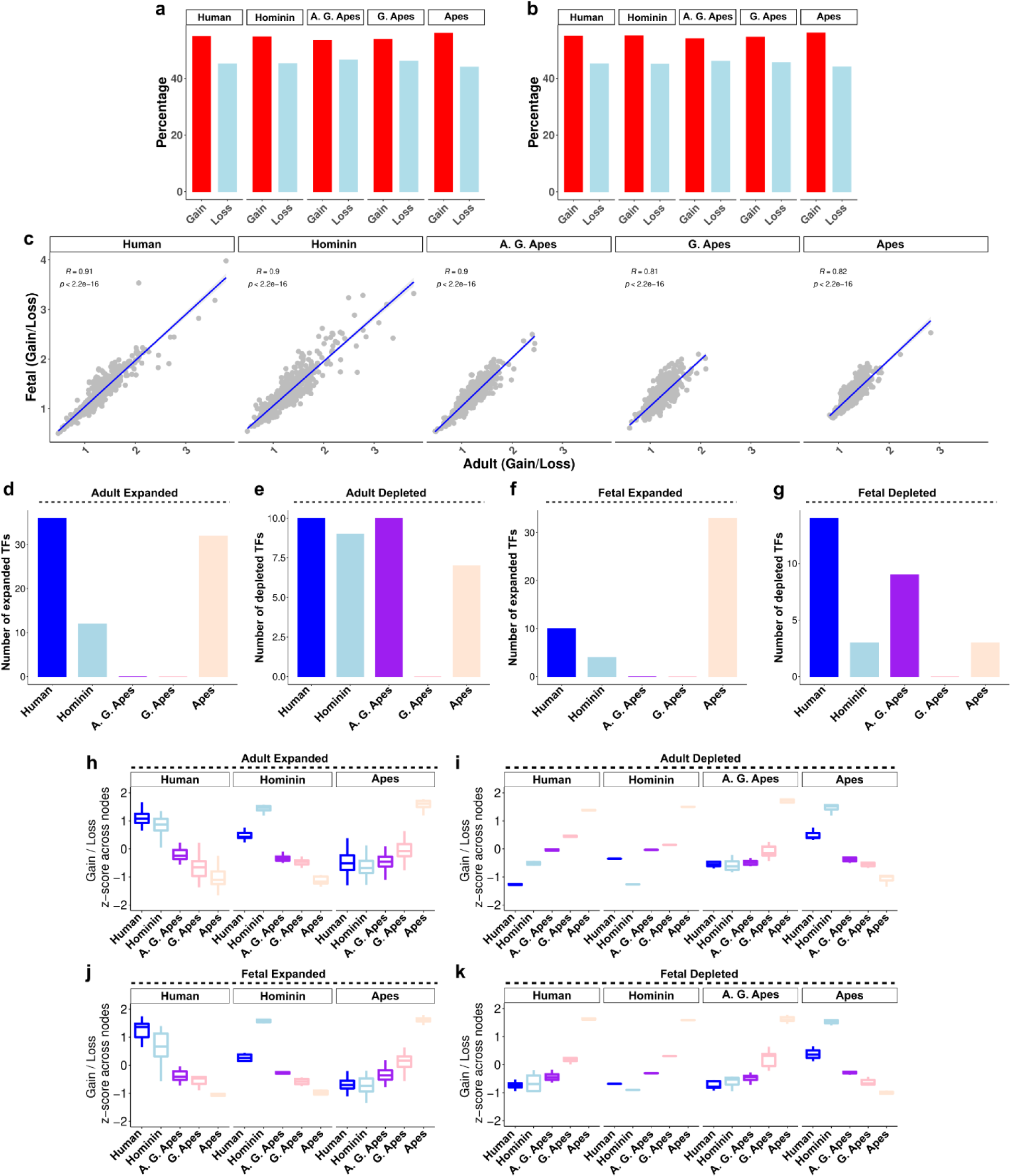
Supplementary analyses on TFBS evolution. **(a-b)** Percentage of TFBS evolution that are gained or lost in each lineage in adult **(a)** or fetal **(b)** datasets. **(c)** Spearman rank correlations of gain / loss ratios between adult and fetal datasets per lineage. **(d-g)** Number of significantly expanded or depleted TFs per lineage in adult **(d-e)** and fetal **(f-g)** datasets. **(h-k)** Z-scores of gain / loss ratios across all lineages per TF group (expanded / depleted in adult / fetal).

**Supplementary Figure 5:**
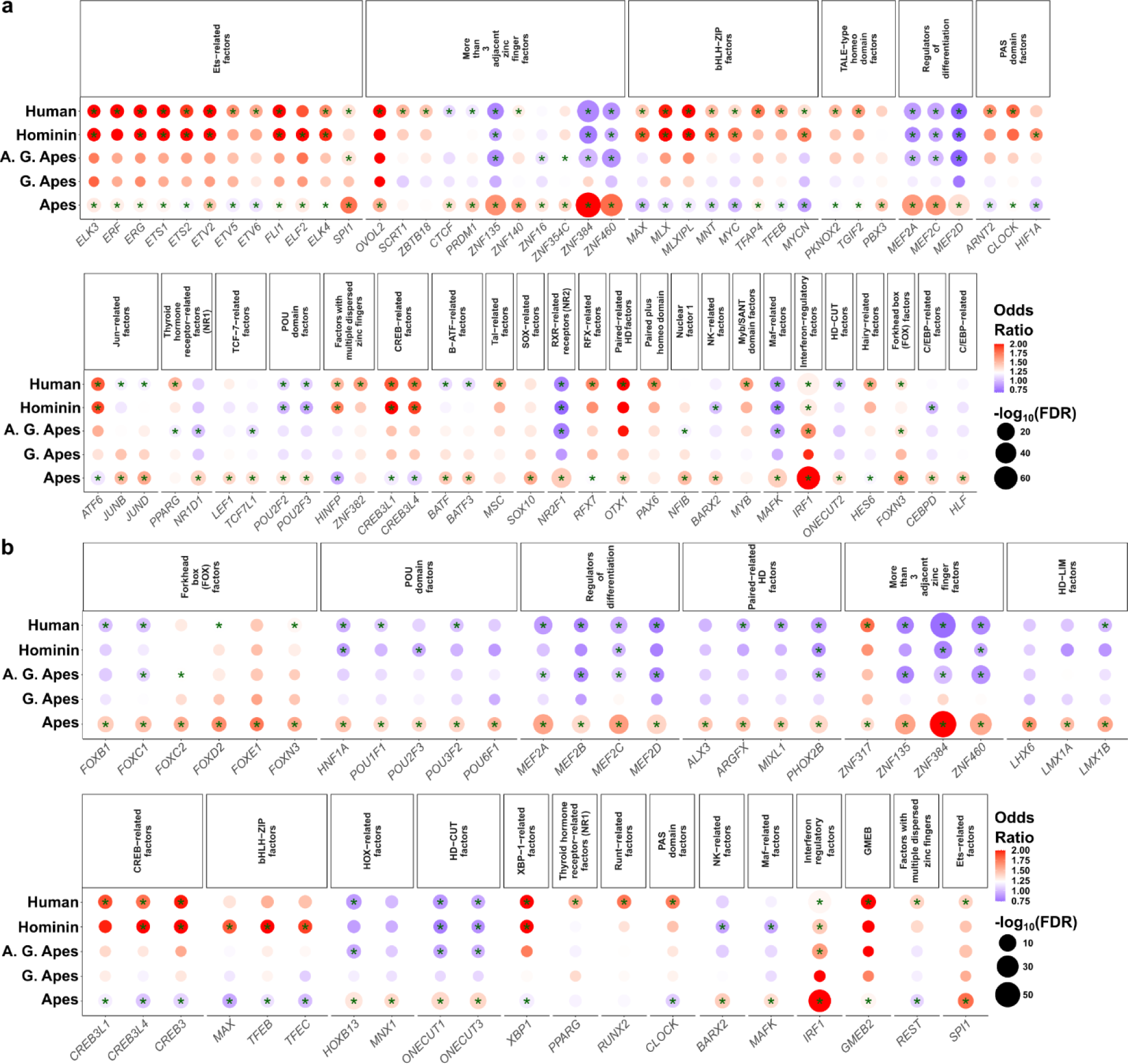
List of all TFBS expansion / depletion. **(a-b)** TFBS expansion / depletion statistics in adult dataset **(a)** and in **(b)** fetal dataset. Asterisk indicates FDR < 0.05. Blue colors indicate depletion, red colors indicate expansion.

**Supplementary Figure 6:**
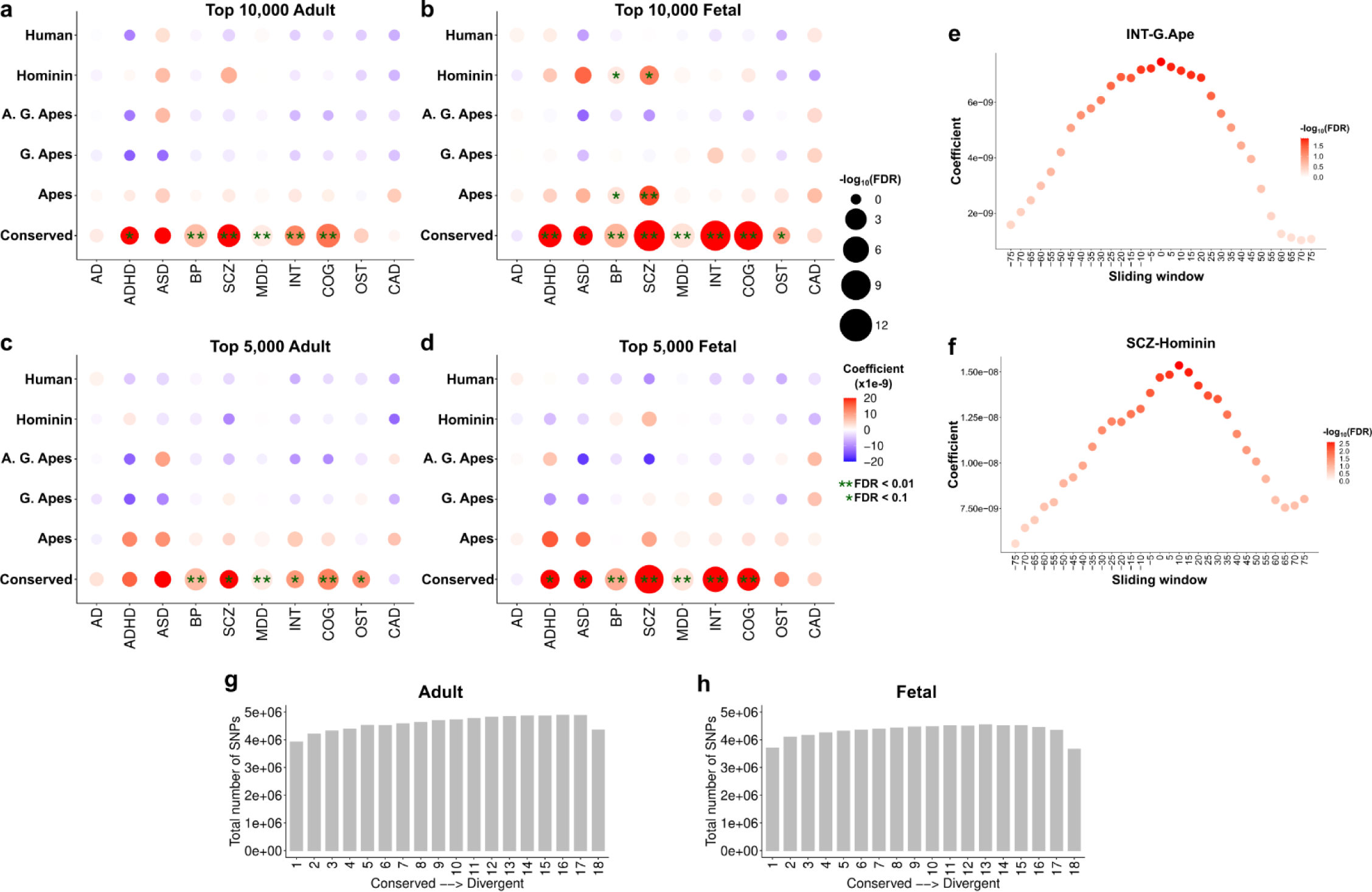
Supplementary results on associations between disease variants and evolutionary divergence. **(a-d)** LDSC regression results between GREs grouped by evolutionary divergence (y-axis) and variants obtained from GWAS studies. **(e-f)** LDSC regressions of SCZ-Hominin and INT-G.Ape association across sliding windows. X-axis denotes the center of the 50kb window (expanded 25kb on each side). **(g-h)** Total number of SNPs mapping to the GREs grouped by conservation in adult **(g)** and fetal **(h)** datasets.

## Supplementary Tables

**Supplementary Table 1: Lineage-specific substitutions in adult human brain epigenome**

This table provides all lineage-specific substitutions for the following lineages: Human, Hominin, A.G. Ape, G. Ape, Ape in the adult dataset.

**Supplementary Table 2: Lineage-specific substitutions in fetal human brain epigenome**

This table provides all lineage-specific substitutions for the following lineages: Human, Hominin, A.G. Ape, G. Ape, Ape in the fetal dataset.

**Supplementary Table 3: GRE groups based on divergence and conservation**

List of all lineage-divergent and conserved GREs for adult and fetal datasets. The table also includes the statistics for the criteria mentioned in the Methods.

**Supplementary Table 4: Cell type marker GREs**

Cell type marker GREs separately for adult and fetal datasets for the broad cell type categories mentioned in text.

**Supplementary Table 5: GRE-Gene links in human brain epigenome**

List of GRE-Gene links based on correlations of GRE accessibility and gene expression in multiomic single-cell adult and fetal brain datasets.

**Supplementary Table 6: Regulation divergent genes (RDGs)**

List of genes linked to significant excess of divergent GREs for each lineage.

**Supplementary Table 7: Gene ontology enrichments**

Statistics of all gene ontology enrichments.

**Supplementary Table 8: TFBS gains and losses**

TFBS gains and losses per lineage and per GRE.

**Supplementary Table 9: TFBS expansions and depletions across lineages**

List of all significant TFBS expansion and deletions across lineages.

**Supplementary Table 10: LDSC statistics**

All LD score regression statistics of the presented results

